# Hypoxia-induced activation of NDR2 underlies brain metastases from Non-Small Cell Lung Cancer

**DOI:** 10.1101/2023.03.20.533395

**Authors:** Jérôme Levallet, Tiphaine Biojout, Céline Bazille, Manon Douyère, Fatéméh Dubois, Dimitri Leite Ferreira, Jasmine Taylor, Sylvain Teulier, Jérôme Toutain, Myriam Bernaudin, Samuel Valable, Emmanuel Bergot, Guénaëlle Levallet

## Abstract

The molecular mechanisms induced by hypoxia are misunderstood in non-small cell lung cancer (NSCLC), and above all the hypoxia and RASSF1A/Hippo signaling relationship.

We confirmed that human NSCLC (n=45) as their brain metastases (BM) counterpart are hypoxic since positive with CAIX-antibody (target gene of Hypoxia-inducible factor (HIF)). A severe and prolonged hypoxia (0.2% O2, 48h) activated YAP (but not TAZ) in Human Bronchial Epithelial Cells (HBEC) lines by downregulating RASSF1A/kinases Hippo (except for NDR2) regardless their promoter methylation status. Subsequently, the NDR2-overactived HBEC cells exacerbated a HIF-1A, YAP and C-Jun-dependent-amoeboid migration, and mainly, support BM formation. Indeed, NDR2 is more expressed in human tumour of metastatic NSCLC than in human localized NSCLC while NDR2 silencing in HBEC lines (by shRNA) prevented the xenograft formation and growth in a lung cancer-derived BM model in mice.

Collectively, our results indicated that NDR2 kinase is over-active in NSCLC by hypoxia and supports BM formation. NDR2 expression is thus a useful biomarker to predict the metastases risk in patients with NSCLC, easily measurable routinely by immunohistochemistry on tumour specimens.

## Introduction

Like other solid tumours, non-small cell lung cancers (NSCLC), the most frequent lung cancers, will experience hypoxia (oxygen deprivation) and subsequent genomic instability, aggressiveness of tumour cells, formation of metastases and resistance to treatment (Chen et al, 2018; Salem et al, 2018; Ziółkowska-Suchanek, 2021). The molecular mechanisms by which hypoxia acts are patchy in lung carcinogenesis. Based on other carcinomas, one can hypothesize that hypoxia could disrupt the RASSF1A (Ras association domain family 1 isoform A)/Hippo signaling pathway (Palakurthy et al, 2009; Yan et al, 2014; Ma et al, 2015; Ma et al, 2016; Thienpont et al, 2016; Wei et al, 2017). In actual fact, YAP is reported to be active in several hypoxic tumours (Zhao et al, 2020). Under hypoxic conditions, the phosphorylation/inactivation of YAP decreases while that of TAZ increases strongly concomitantly as the total form of TAZ in ovarian cancer (Yan et al, 2014). Hypoxia induces YAP interaction with hypoxia-inducible factor-1α (HIF-1A) and its nuclear translocation to promote pancreatic ductal adenocarcinoma invasion via epithelial– mesenchymal transition allowing YAP to be an independent prognostic predictor of survival of these patients (Wei et al, 2017). In line, YAP was shown to bind to the HIF-1A and sustains HIF-1A protein stability to promote hepatocellular carcinoma cell glycolysis under hypoxic stress (Zhang et al, 2018). In addition, another factor in hypoxia belonging to the HIF family, HIF-2A, plays a role in the progression of colon cancer by activating YAP and consequently the expression of these target genes (Ma et al. 2017).

Regarding NSCLC, this hypothesis according to which hypoxia could disrupt the RASSF1A (Ras association domain family 1 isoform A)/Hippo signaling pathway, is of particular interest since Hippo pathway is already known to be altered following the loss of expression of RASSF1A, which occurs in 25% of patient with NSCLC (De Fraipont et al, 2012), leading to aberrant activation of both the Hippo kinase, NDR2 and the Hippo effector, YAP (Keller et al, 2019) and supporting the subsequent initiation and dissemination of NSCLC (Dubois et al. 2016; Keller et al, 2019). To date, only work by Dabral and co-workers reported the role of RASSF1A-HIF-1A loop, in a subset of NSCLC still expressing RASSF1A and the primary cancer cells isolated from the same tumours, independent of Hippo signaling (Dabral et al, 2019). Here, we thus aimed to decipher the relationship between HIF-1A/YAP/TAZ in a context of presence or absence of RASSF1A in Human Bronchial Epithelial Cells (HBEC) lines grown under severe (0.2% O_2_) and prolonged (48h) hypoxia that is, conditions such as those present in the core/bulk of lung tumour, the median tumour pO2 ranged from 0.7 to 46 mm Hg (median,16.6) (Le et al, 2006, McKeown 2014), corresponding to 0.09 to 6.09% O_2_ (median, 2.2%) (Carreau et al, 2011), to identify new tools to establish tumour metastatic score.

## Methods and Protocols

### Patients

We selected a retrospective population of 45 patients operated on a non-metastatic NSCLC (n=25) or metastatic NSCLC (n=20) for whom both the primitive tumour and the brain metastasis (BM) were available, at Caen University Hospital between December 2009 and December 2019. Among the 25 patients with localized NSCLC, 17 were men and 8 were women with an average age of 71 years [54 – 86 years]. Among the 20 patients with metastatic NSCLC, 15 were men and 5 were women with an average age of 67 years [40 – 82 years]. All tumour specimens were reviewed by an experienced pathologist (CB) on slides stained with hemalum-eosin-saffron: the pulmonary origin of BM was certified by the presence of a positive immunostaining for thyroid transcription factor-1, cytokeratin 7 and/or P40 positive immunostaining. As required by French laws, all patients provided informed consent, and the study was approved by the institutional ethics committee (North-West-Committee-for-Persons-Protection-III N°DC-2008-588).

### Mice Brain Metastasis (BM) model

All animal investigations were performed under the current European directive (2010/63/EU) following ARRIVE guidelines. This study was undertaken in the housing and laboratories #F14118001/#G14118001 and with the permission of the regional committee on animal ethics (C2EA-54 CENOMEXA, project #23280). Nude athymic mice (20-25g, 8 weeks, male) were maintained in specific pathogen free housing. Mice were manipulated under general anesthesia (5% isoflurane for induction, 2% for maintenance in a 1L/min of 70%N_2_O/30%O_2_). Body temperature was monitored and maintained at 37.5±0.5°C throughout the experiments. Mice were placed in a stereotactic head holder and a scalp incision was performed along the sagittal suture. A549 or H2030-BrM3 cells (10∧5 cells in 3 μl-PBS supplemented by glutamine (2 mM)) were injected over 5 min (0.6µl/min) *via* a fine needle (30G) connected to a Hamilton syringe. The injection sites were the right caudate putamen at a depth of 4 mm and lateralization on the right of 2.5 mm. Animals were followed twice a week by anatomical MRI over a 21 days period to follow BM development.

### Acquisition of MRI images and MRI sequence analysis

The development of the lesions was monitored twice a week using MRI (magnetic resonance imaging) on a 7 Tesla magnet (Pharmascan, Bruker). All experiments were performed under isoflurane anesthesia: 5% and during induction and 2.5% during the procedure in a 1L/min mixture N20 and 02 (70% and 30%). The mouse is placed in a cradle allowing the head to be held by ear and tooth bars. Breathing is monitored in real time using a pressure balloon under the abdomen.

Fast imaging, FLASH sequence (Fast Low Angle Shot); TR/TEeff: 100/4 msec; resolution 0.39×0.39×3 mm3, acquisition time = 12 sec), was out to verify the positioning of the animal and allow acquisition adjustments. T2-weighted imaging by rapid spin echo or RARE8 (Rapid Acquisition Relaxation Enhanced 8) sequence was then acquired with the following parameters: TR/TEeff = 5000/65 ms, number of repetitions = 1, spatial resolution = 0.078 x0.078, 26 slices 0.5 mm, acquisition time = 2 minutes.

The analyzes of the MRI sequences are carried out to define the different types of lesions and locations to measure the volume of lesions at the different post-injection times. Tumour delineation was performed manually using ImageJ software (NIH, Wayne Rasband, USA) on all adjacent T2w slices and tumour volume was achieved by multiplication of the sum of contiguous tumour surface areas with the slice thickness.

### Cell culture transfection and treatment

Immortalized human bronchial epithelial HBEC-3 was kindly provided by Dr. Michael White (UT Southwestern Medical Center, Dallas, TX, USA), as previously described (Dubois et al, 2016). HBEC-3 cells were grown in keratinocyte serum-free medium (KFSM) supplemented with 0.2ng/mL recombinant Epidermal Growth Factor (EGFr) and 25μg/mL bovine pituitary extracts (BPE) (Thermo Fisher Scientific, Rockford, IL, USA). BEAS-2B, A549, H1299 and H1915 were purchased from the American Type Culture Collection (ATCC). The human H2030-Br3M adenocarcinomas cells (KRAS^G12C^ mutated from MSKCC, Dr Joan Massagué) that preferentially metastasizes to the brain were used. BEAS-2B, A549, H1299, H1915 and H2030-BrM3 were grown in Dulbecco’s Modified Eagle Medium (DMEM) supplemented with 10% (vol/vol) heat-inactivated fetal bovine serum. Both KSFM and DMEM mediums were complemented by 100U/mL penicillin, 100μg/mL streptomycin, and 2mM l-glutamine (Gibco, Life Technologies, Grand Island, NY, USA). The cultures were incubated at 37 °C in a humidified atmosphere with 5% CO2.

RNAi oligonucleotides (Eurogentec®) sequences are detailed in Table S1. The mock siRNA was employed as the non-silencing negative control (Dharmacon, Thermo Scientific, Pittsburgh, PA, USA). Plasmids encoding wild-type RASSF1A (pcDNA3-RASSF1A) and control mimic (Addgene, Cambridge, MA, USA) have been described previously in the supplementary data section (Dubois et al, 2016). The introduction of siRNA and plasmids was performed using Lipofectamine RNAiMax (Invitrogen, Carlsbad, CA, USA) in accordance with the manufacturer’s instructions at 30% (siRNA) and 70% (plasmids) of cell confluence, respectively.

For culture under hypoxia at 0.2% oxygen, the cells were kept in a hermetic hypoxia workstation (INVIVO2, Ruskinn, ABE) with an atmosphere humidified with 0.2% O_2_, 95% nitrogen and 5% CO_2_ with a temperature of 37 ° C.

For the pharmacological inhibition of c-jun, the cells were treated with SP600125 (1 μM) (Selleckchem, Houton, TX, USA).

### Preparation of RNA and RT-PCR

The extraction of total RNA from cells was carried out using the illustra RNAspin mini® column (GE Healthcare, Bio-Sciences, Pittsburgh, PA, USA), according to the manufacturer’s instructions. Total RNA was treated with DNAse I (Invitrogen, Carlsbad, CA, USA) to discard contaminating genomic DNA. The RNA concentrations were determined using spectrophotometer Nanodrop® 2000c. Total RNA (250 ng) was reverse-transcribed with random primers and 200 IU M-MLV reverse transcriptase at 37 °C for 90 min, followed by 5 min of dissociation at 70 °C with Mastercycler Eppendorf®. The resulting cDNAs were diluted (1/10) and used as templates. Polymerase chain reaction (PCR) was performed in a Mx3005P QPCR system (Agilent Technology) with 5 pmol of each primer set (Table S2) and iQTM SYBR Green Supermix (Bio-Rad, Hercules, CA, USA) as follows: 95 °C for 5 min, followed by 40 cycles at 95 °C for 1 min, and annealing/extension at 60 °C for 60 s. S16 was used as an internal control. Positive standards and reaction mixtures lacking the reverse transcriptase were employed routinely as controls for each RNA sample. Relative quantification was calculated using the ΔΔCt method.

### Preparation of DNA and methylation-specific PCR assay

DNA samples obtained from cells using the QIAamp DNA Tissue kit (Qiagen). Genomic DNA bisulfite modification was performed using the Epitect kit (Qiagen), according the manufacturer’s instructions and as previously described (Levallet et al, 2019). Polymerase chain reaction (PCR) was conducted with specific primers for either the methylated or unmethylated alleles in standard conditions for the following genes encoding proteins of the Hippo pathway or RASSF superfamily: RASSF1A, MST1, MST2, LATS1, LATS2, NDR1, NDR2 and ANKRD1 (Table S3).

### Antibodies, Immunofluorescence (IF), immunohistochemistry (IHC), immunoblotting and image analysis

The antibodies used in this work are listed in Table S4.

For IF studies, cells were seeded on coverslips in 24-well tissue culture trays at a density of 2 × 10^4^. Following 48 hours, cells were washed with PBS and fixed with 4% paraformaldehyde for 20 minutes at 37 °C. The cells were then permeabilized with frozen methanol for 10 minutes and blocked with 4% bovine serum albumin (BSA) in phosphate-buffered saline (PBS) for 1 hour and stained with primary antibodies at 4 °C overnight. After being washed with PBS, cells were stained with either Alexa-488-conjugated or Alexa-555-conjugated secondary antibodies (Molecular Probes, Invitrogen, Eugene, OR, USA) for 1 hour at room temperature (RT) and with DAPI (4,6 diamidino-2-phenylindole) (Santa Cruz Biotechnology, Dallas, TX, USA). Digital pictures were captured using a high-throughput confocal microscopy (FluoView FV1000, Olympus).

For IHC studies, tumour paraffin-embedded blocks were processed as previously described (Levallet et al, 2012). Slides were pretreated with 0.01 mol/L citrate buffer (pH 6; Dako) or EDTA solution (pH 9; Dako) for 20 minutes at 100°C, then immunostainings were performed with automated immunohistochemical stainer (Dako). Slides were successively incubated at room temperature in 3% H2O2 for 5 minutes, then with primary monoclonal antibody diluted at 1:50 to 1:1000 for 60 minutes at room temperature. Finally, antibody fixation was revealed by the Novolink System (Leica). The staining intensities were evaluated in a blinded manner at 40x magnification and were scored using marker-specific 0–3 scales (0: negative, 1: weak, 2: moderate, and 3: strong). An overall IHC composite score was calculated based on the sum of the staining intensity (0–3) multiplied by the distribution (0%– 100%) from all parts of the slide, thereby providing an H-score between 0 and 300.

For immunoblotting, whole-cell protein extracts were prepared as previously described (Dubois et al, 2016), and proteins were detected by immunoblotting with the primary antibody diluted to 1:1000 in Tween (0.1%)-TBS buffer and horseradish peroxidase (HRP)-conjugated secondary antibody, then revealed by enhanced chemiluminescence using the ECL kit (Promega™). Densitometry results of western blot were analyzed with Image J software. The signal intensity of each band was normalized with GAPDH densitometry values.

### BrdU incorporation assay

BrdU incorporation assay kit was used in accordance with the manufacturer’s instructions (1/500 dilution) (cat. no. 2750, Millipore, Billerica, MA, USA) and detected by adding a peroxidase substrate. Spectrophotometric detection was performed at 450nm wavelength.

### Apoptosis measurement

DNA fragmentation and caspase-3/7 activation were assayed using the Cell Death Detection ELISA Plus Kit (Roche, USA) and the Caspase-Glo 3/7 Luminescence Assay (Promega Corp., Madison, WI, USA), respectively, according to the manufacturer’s instructions.

### Wound-healing assay

Transfected cells were seeded on collagen IV-coated plates (BD BioCoat Matrigel® Invasion Chamber, 8.0-μm pore size for 24-well plates) (BD Biosciences, Bedford, MA, USA), grown to confluence, and pretreated with mitomycin C (1μg/mL) over 12 hours before we created an artificial “wound” by scratching with a P-200 pipette tip (point “0 h”). The cells were then allowed to close the wound and were photographed at six hours (point “6h”). The wound was photographed under a microscope (x10). Wound healing was measured as μm/hour by calculating the reduction in the wound’s width between 0 and 6 hours.

### 3D migration assay

The total of 25 × 10^3^ cells in 250μL serum-free medium were placed in the upper chambers of 24-well Transwell plates containing a cell culture inserted with 8 μm pore size (Greiner Bio-One, Courtaboeuf, France). The lower chamber was filled with 700 μL complete media. After 48 hours of incubation, the non-migrating cells on the top were removed using cotton swabs, and the migrating cells on bottom surface of the filter were stained using crystal violet. Quantification of the migration and invasion assay was performed by counting the cells on the filter’s lower surface under an inverted microscope at x20 magnification. Assays were performed in triplicate, with data presented as the average number of invading cells.

### Statistical analysis

Data were expressed as means ± SEM of experiments, independently conducted three times. All statistical analysis was performed using GraphPad Prism 4, a GraphPad Software program (San Diego, CA, USA). The data were analyzed using a two-tailed Student’s t-test for single comparison or one-way ANOVA followed by Dunnett’s multiple comparison analysis. Differences were considered significant at the p < 0.05 level.

## Results

### Human primitive NSCLC as their brain metastases are hypoxic

Brain metastases from patients with NSCLC are hypoxic tumours as shown by positive immunostaining with carbonic anhydrase 9 (CAIX), a transcriptional target of HIF-1A (Corroyer-Dulmont et al, 2021). Here, we further report that the CAIX H-Score is similar between primary tumours of patients with localized NSCLC (62.4 ± 12.3) and those with metastatic NSCLC (71.2 ± 19.7), and comparable between primary tumours and brain metastasis (65.4 ± 17.1, n=20) from the same patients (Figure S1).

### Severe and prolonged hypoxia (0.2% O_2_, 48h) activates YAP but not TAZ in HBEC lines

We grew HBEC lines in severe (0.2% O_2_) and prolonged (up to 48h) hypoxia (Figure S2). The nuclear accumulation of HIF-1A confirmed the activation of the hypoxia-dependent pathway after 48h of culture in hypoxia at 0.2% O_2_ (Figure S2A and S2B for respectively HBEC-3 and A549 cells). Cell labeling by crystal violet reveals that HBEC-3 (Figure S2C) or A549 cells (Figure S2D) are significantly less dense for the cells cultured in hypoxia 0.2% O_2_ for 24h or 48h, compared to the normoxia condition. Caspase3/7 activity (apoptosis) significantly increases in HBEC-3 (Figure S2E) or A549 (Figure S2F) in severe (0.2% O_2_) and prolonged (up to 48h) hypoxia. However, HBEC-3 cells still incorporate BrdU between 24 and 48 hours (Figure S2G) as A549 cells (Figure S2H). We confirm thus that HBEC-3 (Figure S2) as the other HBEC lines used to survive to such hypoxia (48h, 0.2% O_2_) (Figure S3).

Next, we analyzed the protein expression of the total and phosphorylated (inactivated) forms of the terminal cofactors of the Hippo pathway, YAP and TAZ, by Western blot in HBEC-3 cells (Figure 1). Hypoxia (0.2% O_2_, 48 h) increases by 3-fold the total form of the YAP protein which accumulates in its active (dephosphorylated) form in HBEC-3 while the expression of TAZ drastically decreases except when the cells lack both RASSF1A and YAP: the silencing of YAP and/or RASSF1A increase in additive way the expression of TAZ expression in both normoxia and hypoxia conditions (Figure 1A). Silencing TAZ or YAP by siRNA confirmed specificity of immunoblot (Figure 1A). We then immunostained the YAP and TAZ proteins to determine their subcellular location (Figure 1B-C) and quantified the expression of two of its target genes (Figure 1D-E). We observe that hypoxia (0.2% O_2_, 48h) significantly increases the YAP nuclear intensity like the RASSF1A loss of expression and that there is no additive effect between these two events (Figure 1B) while TAZ nuclear signal drastically decreases under such growth culture (Figure 1C). The quantification of *CTGF* and *ANKRD1* mRNAs expression, some of the target genes of YAP (Meng et al, 2016; Jiménez et al, 2017) confirms that YAP is active when the HBEC-3 cells are cultured in hypoxia (0.2% O_2_, 48h) (Figure 1D). The expression of the CTGF mRNA is significantly reduced by half when the HBEC-3 cells are cultured in hypoxia, while the expression of the *ANKRD1* mRNA is significantly increased. The loss of expression of RASSF1A, which causes the nuclear translocation of active YAP in HBEC-3 cells (Dubois et al, 2016), significantly increases the expression of these two target genes of YAP as the cells are cultured in normoxia or in hypoxia (Figure 1D-E).

**Figure 1:**
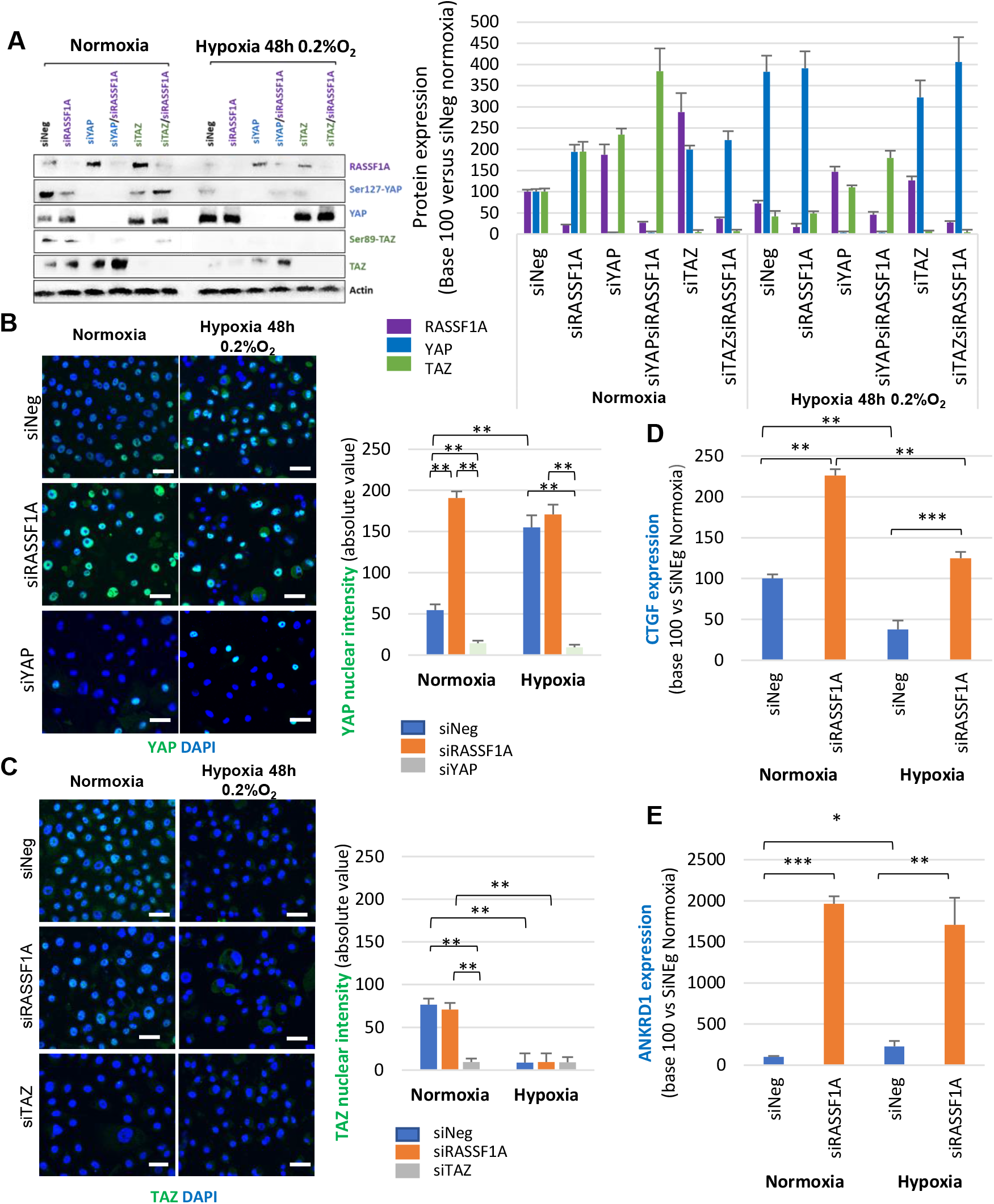
Severe and prolonged hypoxia activates YAP and downregulates TAZ in HBEC-3 cells. **A**) Western blot of members of the Hippo pathway on HBEC-3, expressing or not RASSF1A, after 48 h of normoxia/hypoxia and associated quantification (right panel). (**B** and **C**) Immunostaining of YAP and TAZ respectively on HBEC-3 expressing or not RASSF1A and YAP or TAZ, after 48h of normoxia/hypoxia and Graphic representation of YAP or TAZ nuclear intenstity. **D** and **E**) Graphic representation of the mRNAs expression of the target genes of YAP, CTGF (**D**) and ANKDR1 (**E**) P-value * P <0.05, ** P <0.01 and *** P <0.001 (SEM n≥3).

To determine whether a severe and prolonged hypoxia (0.2% O_2_ for 48h) modulate the activity of YAP and/or TAZ in NSCLC lines, we analyzed the protein expression of their total and phosphorylated (inactivated) forms by Western Blot (Figure S4A). The phosphorylated form of TAZ (p-TAZ) as TAZ saw its expression significantly decrease in all lines (from -40 to -80%) with the exception of line H1299 for which the decrease of pTAZ was insignificant (-19%) (Figure S4B-C). Unlike its paralog, the expression of YAP is less affected by a severe and prolonged hypoxia of HBECs and seems rather stabilized in all cell lines (Figure S4D). For the phosphorylated form of YAP (p-YAP), we do not report a significant difference in phosphorylation between normoxia and hypoxia (Figure S4E). We next confirmed that YAP activation induced by a severe and prolonged hypoxia was not unique to HBEC-3 cells by performing the same experiments with others HBEC cells. The nuclear intensity of YAP in hypoxic condition seems greater for the A549 lines with an increase of 45.0 ± 14.3% in intensity and 69.9 ± 25.0 % for the H1299 cell line (Figure S4F), which would accumulate in its active form as evidenced by the significant increase in *CTGF* expression and *ANKRD1* quantified by qRT-PCR (Figure S4G). We note that an increase in YAP nuclear intensity is also found in BEAS-2B but not significantly, while in the H1915 and H2030-BrM3 cell lines no variation of YAP localization was found (Figure S4F). Notably, we observe a variation in the expression of the different target genes (expression of *CTGF* vs *ANKDR1* in H1299 cells) inside a cell line as well as between the cell lines (*ANKDR1* in A549 vs H1299) (Figure S4G) suggesting that the hypoxia-induced transcriptional regulation depends on the cell line studied as well as on the target gene. Regarding TAZ, we observe a drop in protein expression and nuclear intensity in the four HBEC lines tested when they are incubated in severe hypoxia and this significantly for all cell lines (Figure S4H). At the same time, in all these cell lines, HIF-1A accumulates in the nucleus of cells when cultured under hypoxia. (Figure S4I).

### Severe and prolonged hypoxia (0.2% O_2_, 48h) downregulated RASSF1A/kinases Hippo in HBEC lines except for NDR2

The activity of YAP is inhibited when YAP is phosphorylated by the kinases of the Hippo pathway, the NDRs, themselves activated by the kinases MST1/2. We therefore verified the effect of hypoxia on the expression of mRNAs of different kinases of the Hippo pathway by qRT-PCR in HBEC-3 (Figure 2). These results show an inhibitory effect of hypoxia on the expression of the messengers of the kinases MST1, LATS1 and NDR1/2 (Figure 2A-D). We also find that the silencing of RASSF1A in HBEC-3 cells is followed by increased expression on NDRs kinases in normoxia for NDR1 or in both normoxia and hypoxia condition for NDR2 (Figure 2C-D). Finally, at the scale protein we confirm that severe and prolonged hypoxia (0.2% O_2_ for 48h) drastically reduces the expression of Hippo kinases MST1, LATS1 and NDR1 while NDR2 is preserved (Figure 2E).

**Figure 2:**
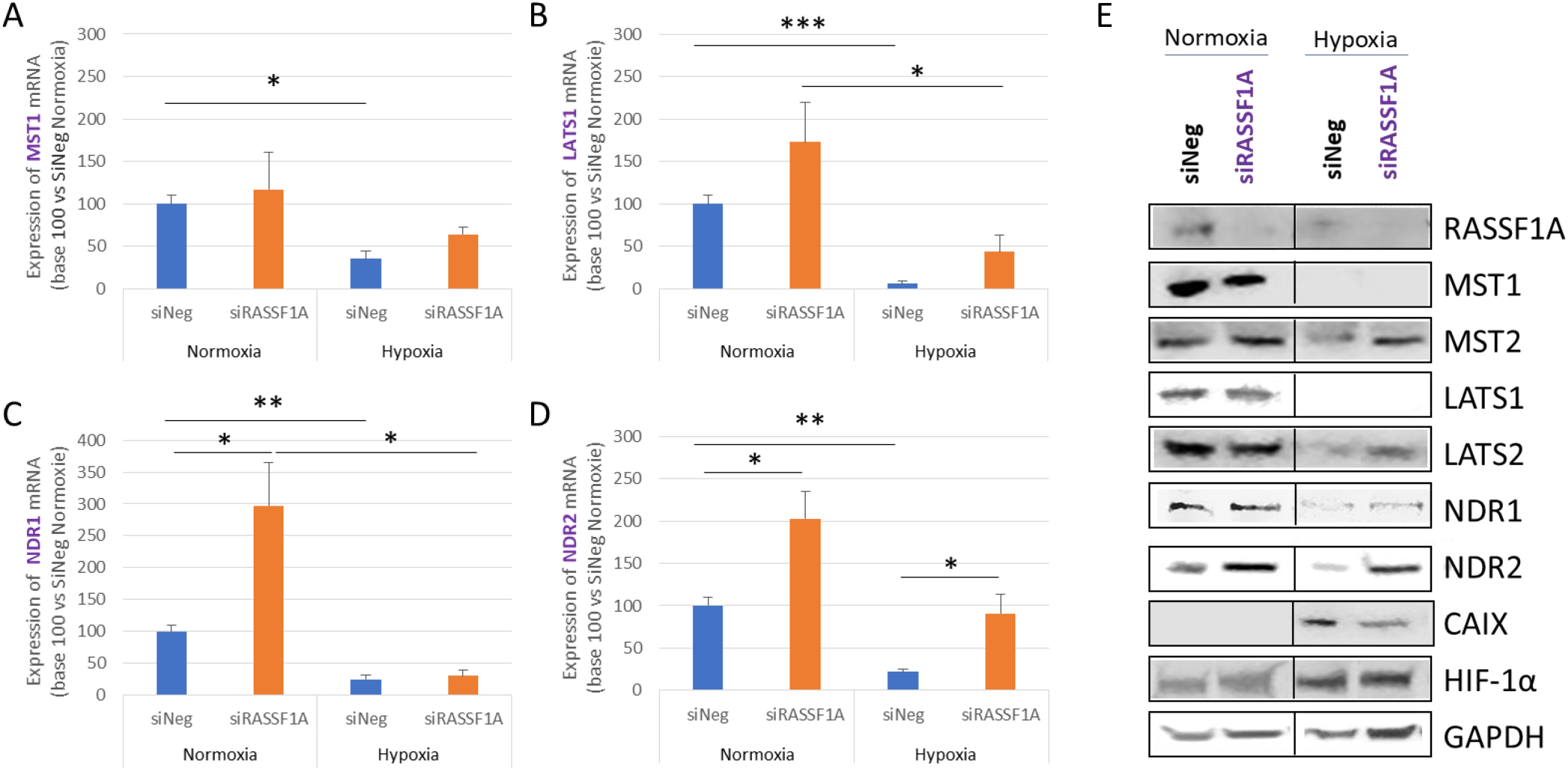
Severe and prolonged hypoxia preserves NDR2 protein expression but decreases expression of other Hippo pathway kinases in HBEC-3 cells. **A-D**) Graphic representation of the mRNAs expression of the kinases of the Hippo pathway MST1, LATS1 and NDR1/2 (A-D) by qPCR on the HBEC-3 cells. P-value * P <0.05, ** P <0.01 and *** P <0.001 (SEM n≥3). **E**) Western blot of Hippo pathway members in HBEC-3 cells, expressing or not RASSF1A, after 48 h of normoxia / hypoxia (representative experiment).

Moreover, as shown in Figure S5, the protein expression of LATS1, LATS2 and NDR1 kinases is significantly reduced (with the exception of LATS1 in A549 cells) in the BEAS-2B, A549, H203-BrM3 lines cultured under hypoxia (Figure S4I). In H1915 cells, a tendency to a decrease expression of LATS1 and LATS2 under hypoxia is also observed but not for NDR1 or NDR2. Conversely, none of these 4 kinases sees its expression significantly influenced by hypoxia in the H1299 line. Remarkably, we can observe that the expression of the NDR2 kinase is not significantly modified by hypoxia, whatever the cell line considered (Figure S5E).

Hypoxia could lead to DNA hypermethylation in some tumours including NSCLC (Thienpont et al, 2016). In order to determine whether the decrease in expression of Hippo kinase, which we report when HBEC-3 cells are cultured in hypoxia, was due to hypermethylation of their promoter, we evaluated the methylation status of these promoters by PCR specific methylation or COBRA PCR (for the *ANKRD1* promoter) (Figure S6). We demonstrate that hypoxia (0.2% O_2_, 48h) has no effect on the hypermethylation (HBEC-3) (Figure S6A) / demethylation (A549) (Figure S6B) of the promoters of the genes encoding the different members of the Hippo pathway or the target gene of YAP, *ANKRD1*, known to be inactivated/hypermethylated in NSCLC (Jiménez et al, 2017) (Figure S6).

Taken together, these results therefore report that the HBECs cultivated in severe and prolonged hypoxia most often exhibit an activation of YAP (but an inactivation of TAZ) consistent with the conservation of the expression of NDR2 and the inactivation of the other kinases of the Hippo pathway.

### Hypoxia exacerbates the ability of NDR2-overactived NSCLC cells to perform a YAP/C-Jun and HIF-1A-dependent amoeboid migration

By assess the cell morphology by co-immunolabeling two elements of the cytoskeleton: actin (green) and tubulin (red) filaments, with a confocal laser scanning microscope, we noticed that the cells of the HBEC-3 placed in hypoxia are individualized and adopted stretched and/or even branched positions (Figure 3A). We also observe that both actin and tubulin are in their polymerized form, as evidenced by the presence of visible microfilament structures in normoxia as under hypoxia. Finally, few figures of apoptosis were observed whether these cells were cultured in normoxia or hypoxia (data not shown).

**Figure 3:**
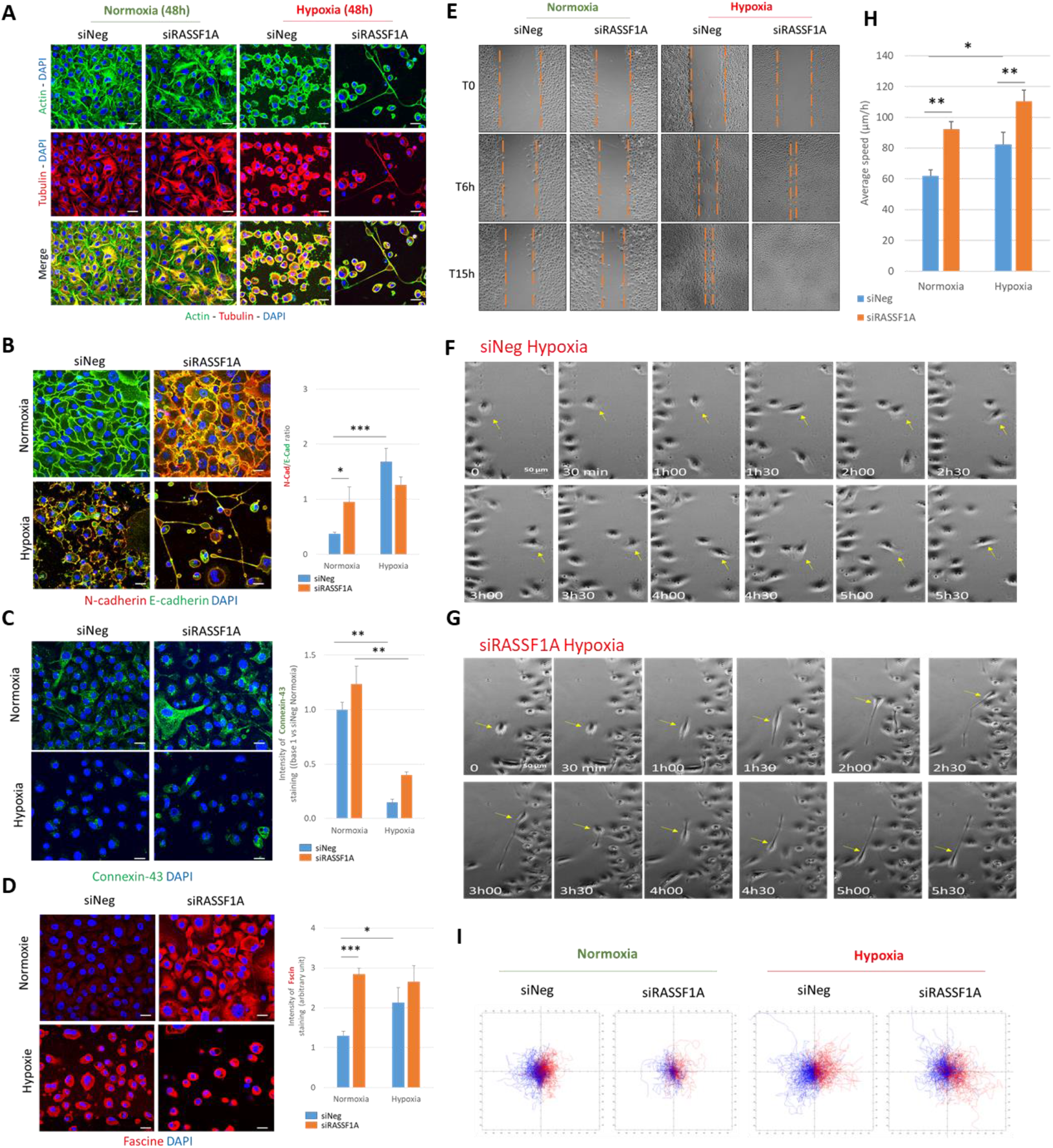
Severe and prolonged (0.2% O_2_, 48h) hypoxia leads to epithelial-mesenchymal transition, cell junctions disrupt, elasticity increase of HBEC-3 cells and thus to increase in individual type migration. **A-D**) Immunostaining on HBEC-3 expressing or not RASSF1A after prolonged culturing (48h) in normoxia/hypoxia showing (**A**) elements of the cytoskeleton, actin (green) and tubulin (red), (**B**) the N -cadherin (in red) and E-cadherin (green) (**C**), connexin 43 (Cx43) (green) (**D**) and fascin (red). (bare Scale 40 µm) (N=3). **E-H**) Illustrations of the wound healing assay with HBEC-3 expressing or not RASSF1A, taken by inverted phase-contrast microscope (x10 magnification) in normoxia/hypoxia, at T0, T6 and T15h after scraping (**E**) zooming in on cell migration for conditions siNeg-hypoxia (**F**) and SiRASSF1A-hyopoxia (**G**). (**H**) The average cell velocity (in µm/h) was measured in normoxia and hypoxia for the control conditions, in the absence of RASSF1A. P-value *P<0.05, **P<0.01 (SEM n≥3). (**I**) Diagram representing the migration and the change of direction of HBEC-3, with in red the cells of the right edge of the wound and in blue those of the left edge (>300 cells) using the MtrackJ® module of the Fiji® software.

We sought to evaluate the effect of this hypoxia on the epithelial-mesenchymal transition, the adherent and communicating junctions and the elasticity of HBEC-3 cells by measuring the expression of E- and N-Cadherins (Ashaie et al, 2016), connexin43 (Kotini et al, 2018) and fascine (Tanaka et al, 2019) (Figure 3B-D). We observe a decrease in the expression of E-cadherin (epithelial marker) and an increase in N-Cadherin (mesenchymal marker) in HBEC-3 cells placed in hypoxia (0.2% O_2_, 48h) compared to cells cultured in normoxia (Figure 3B). We also observe a decrease of these proteins at the membrane surface when cells are cultured in hypoxia (0.2% O_2_, 48h) compared to cells in normoxia (Figure 3C). In HBEC-3 cells cultured in hypoxia (0.2% O_2_, 48h), we further observe the loss of connexin 43 expression by a factor of 5 but a cytoplasmic increase of fascin expression (Figure 3D). In HBEC-3 cells depleted to RASSF1A expression, loss of connexin 43 is still evidence in hypoxia but not the concomitant increase of fascin (Figure 3D).

Migration of HBEC-3 cells is assessed *via* a wound healing test under normoxia or in hypoxia (0.2% O_2_) condition (Figure 3E). This real-time monitoring of wound filling reveals that the “control” HBEC-3 cells, grown in hypoxia (0.2% O_2_, 48h), adopt an individual migration mode while their migration is collective in normoxia (Figure 3E). This migration is of amoeboid type for the “control” cells (Figure 3F) while mesenchymal for RASSF1A-depletd HBEC-3 cells (Figure 3G).

Since the type of migration (individual *versus* collective) influences the speed of cell movement (Lintz et al, 2017), we measured the average speed of cell movement using the TrackMate module of the Fiji® software over more than 500 cells per film. These analyzes demonstrate that hypoxia (0.2% O_2_, 48h), like the inactivation of RASSF1A, significantly increases the migration speed of HBEC-3 cells without additive effect (Figure 3H). Finally, we observe that the HBEC-3 cells cultured in hypoxia migrated faster without however reaching the other bank. We evaluate the route of these cells using the MtrackJ® module of the Fiji® software, and show that compared to HBEC-3 cells in normoxia, HBEC-3 cells in hypoxia move randomly above all when depleted for RASSF1A (Figure 3I).

Hypoxia and inactivation of RASSF1A have a similar effect on increasing the velocity of HBEC-3 cells. Additionally, inactivation of YAP partly prevents the gain in migration velocity induced by inactivation of RASSF1A when HBEC-3 cells are cultured in hypoxia at 0.2% O2. Thus, we wondered what part HIF1, described to induce cell movements *via* RhoA in particular (Tátrai et al, 2017), took in this 2D migration increase. *HIF-1A* mRNA expression was assessed by qRT-PCR on HBEC-3 line at 48h normoxia/hypoxia (Figure 4A). The results show that the inhibition of the expression of RASSF1A increases by more than three-fold the *HIF-1A* mRNA under conditions of normoxia while *HIF1-A* mRNA expression was significantly reduced by a factor of four in HBEC-3 cells grown under hypoxic condition (0.2% O_2_, 48h).

**Figure 4:**
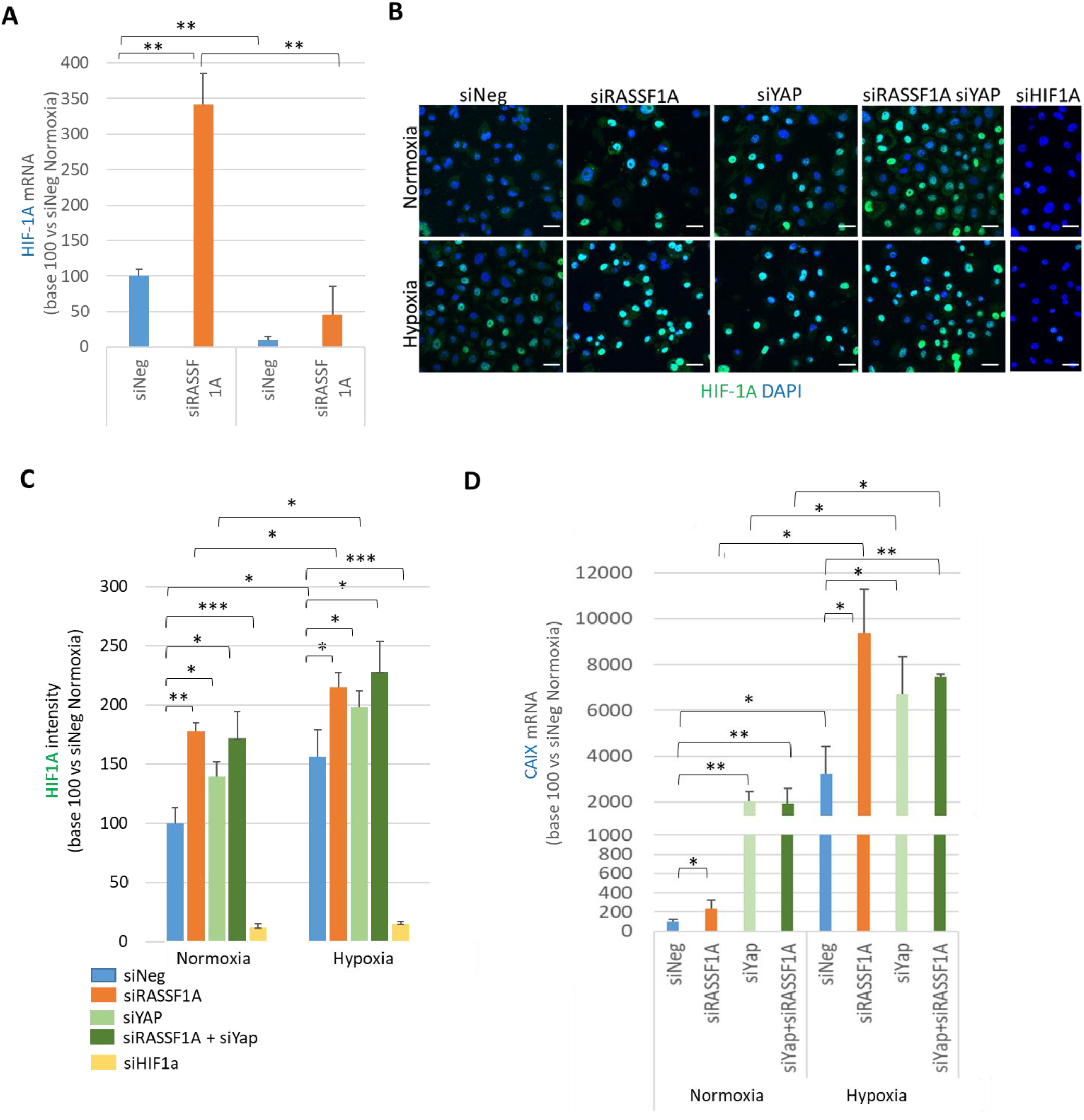
HIF-1A is activated by loss of RASSF1A and/or YAP expression in HBEC-3 cells cultured in hypoxia (0.2% O_2_, 48h). (**A**) Expression of HIF1A mRNA measured by qPCR in HBEC-3 after 48h of normoxia/hypoxia. Illustrations (**B**) and quantification (**C**) of HIF-1A on HBEC-3 expressing or not RASSF1A, YAP or HIF-1A (40 μm scale). (**D**) Expression of *CAIX* mRNA in HBEC-3 after 48 h of normoxia/hypoxia. Values are means ± SEM expressed in base 100 over siNeg in Normoxia. P-value *p<0.05, **p<0.01 and ***p<0.001.

We then evaluated the effects of hypoxia and/or loss of RASSF1A and/or YAP expression on HIF-1A protein expression in immunofluorescence (Figure 4B). We observe that the presence of HIF-1A in the nucleus of HBEC-3 cells at high cell density whether cultured in normoxia or hypoxia and that the loss of RASSF1A and/or YAP enhances the nuclear intensity of HIF-1A (Figure 4C). We quantified the expression of one of its target genes, carbonic anhydrase IX (CAIX, Wykoff et al, 2000). The quantification of *CAIX* mRNA, evaluated by qRT-PCR, shows that hypoxia induces a significant increase in the expression of *CAIX* mRNA in HBEC-3 cells after 48 hours of hypoxia (Figure 4D). This expression is even higher when RASSF1A and/or YAP are inactivated in cells whether in normoxia or hypoxia (Figure 4D) which is in agreement with the immunostaining data for HIF-1A (Figure 4C). These data demonstrate that hypoxia and/or loss of RASSF1A and/or YAP expression activate the hypoxia factor HIF-1A.

We next evaluated the involvement of c-Jun in the positive effect of hypoxia on NDR and YAP activity by measuring the phosphorylation of c-Jun by western blot and the subsequent consequences on migration in presence or absence of an inhibitor of JNK, the SP600125. Indeed, 1) c-jun is involved in cell motility, cell invasion and TEM (Grose, 2003, Whang et al, 2017, Lin et al, 2018), 2) c-Jun and HIF-1 functionally cooperate in hypoxia-induced gene transcription (Alfranca et al, 2002), 3) c-Jun protects HIF-1A from degradation (Yu et al, 2009) while, 4) prolonged or chronic hypoxia stimulates expression of the stress-inducible transcription factor gene *c -jun* (Laredoute et al, 2002), 5) RASSF1A is a repressor of c-jun in lung cells (Whang et al, 2005) and finally, 6) c-jun is a transcription factor for both YAP-1 (https://www.genecards.org/cgi-bin/carddisp.pl?gene=YAP1) and ARNT (https://www.genecards.org/cgi-bin/carddisp.pl?gene=ARNT).

We show that, in normoxia the loss of RASSF1A increase phosphorylation of c-Jun in the HBEC-3 cells. In contrast silencing of YAP (by siYAP) decreases p-c-Jun/c-Jun ratio and abrogates effect of the loss of RASSF1A (Figure 5A). In hypoxia condition, none of these effects had been observed as well as, when cells were pre-treated with JNK inhibitor which significantly reduced c-Jun phosphorylation. We show that, in normoxia, the loss of RASSF1A increase phosphorylation of c-Jun in the HBEC-3 cells (Figure 5A). In contrast silencing of YAP (by siYAP) decreases this phosphorylation and abrogates effect of the loss of RASSF1A. In parallel, we observed that loss of RASSF1A induces significant increase of HBEC-3 cells velocity (Figure 5B) and 3D migration (Figure 5C) while hypoxia leads to opposite effects on HBEC-3 cells velocity (increase) and migration (decrease). Furthermore, YAP mediates the increase of velocity and migration induced by the loss of RASSF1A expression in normoxia while in hypoxia the increase of cell velocity and the decrease of 3D migration is always observed in absence of RASSF1A and/or YAP expression. The JNK inhibitor abrogates (Figure 5B) or reduces (Figure 5C) effects of loss of RASSF1A expression or hypoxia on HBEC-3 cell velocity and 3D migration respectively.

**Figure 5:**
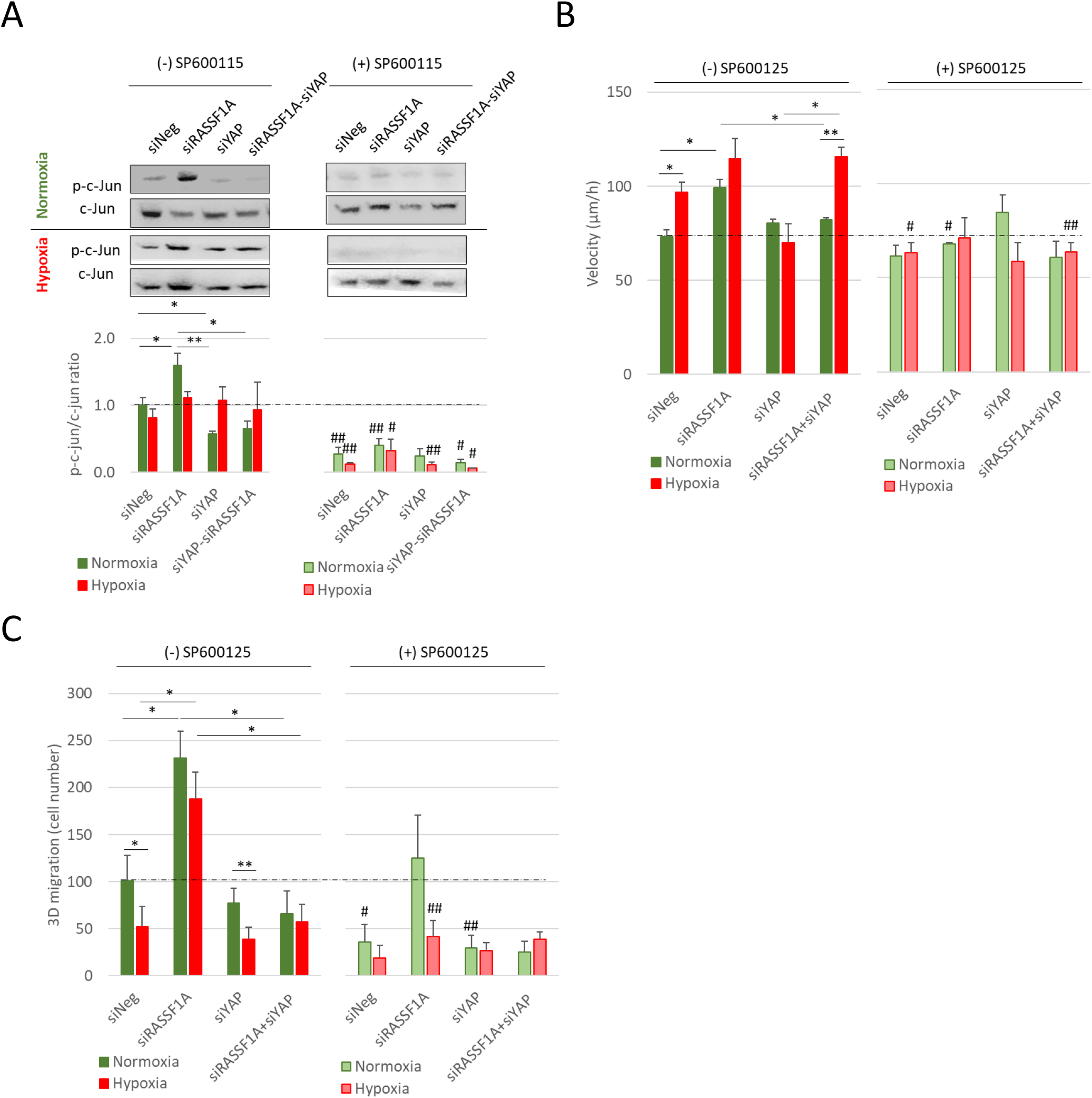
Hypoxia exacerbates the ability of HBEC cells with overactive NDR2 to migrate in a YAP and C-Jun-dependent mechanism. **A**) Expression of p-c-Jun, c-Jun and proteins evaluated by western blot in HBEC-3 cells transfected with siNeg, siRASSF1A, siYAP or both and cultivated for 48 hours in normoxia or hypoxia (0.2% O_2_). Upper panels are representative experiments and lower panel is densitometric analysis of p-c-Jun/c-Jun ratio expressed in base 1 using siNeg in normoxia condition. (**B**) Cells velocity and 3D migration (**C**) of HBEC-3 cells transfected with siNeg, siRASSF1A, siYAP or both and cultivated for 48h in normoxia or hypoxia (0.2% O^2^). The values are the mean +/-SEM of 3 independent determinations. ANOVA was followed by a post-hoc Dunnett test, *: p <0.05, **: p <0.01 or #: p<0.05, ##: p <using t-test by comparing (-) SP600125 vs (+) SP600125 in the same culture conditions.

### NDR2 silencing strongly inhibits the xenograft formation and growth in a mice BM model

We use then a lung cancer-derived brain metastases (BM) model in mice, and inoculated H2030-BrM3 cells (shControl, shNDR1 or shNDR2), in the right caudate putamen of Nude athymic mice (N=10 per condition). Stability of the NDR1 and NDR2 silencing were checked out over successive cell passage and controlled prior to inoculation (Figure 6A-C). Eighteen days after cells inoculation, BM were observed in striatum of 6 and 7 animals over 10 in shControl and shNDR1 experimental group respectively while none of mice receiving NDR2-invalidated H2030-BrM3 cells have developed BM (Figure 6D). At day 24, brain metastasis volume reached 36.22 ± 5.8 mm^3^ (for 7 animals/10) in shNDR1 group and 18.73 ± 4.5 mm^3^ (for 7/10) for the shControl one. In shNDR2 group, 6 animals started to developed BM at day 24, with significantly lower average volume (reaching 2.83 ± 0.9 mm^3^) compared to shControl and shNDR1 experimental group. One representative MRI image of BM from each group is shown in right panel (Figure 6D) and the images for all mice brain are in Figure S7. Of note, comparable responses were obtained following the xenograft obtained from cells A549, expressing or not NDR1 or NDR2 (not shown data). One could also note that the tumor structure looked similar, ie., slightly hemorrhagic as they appear dark in T2w images for the three group of mice.

**Figure 6:**
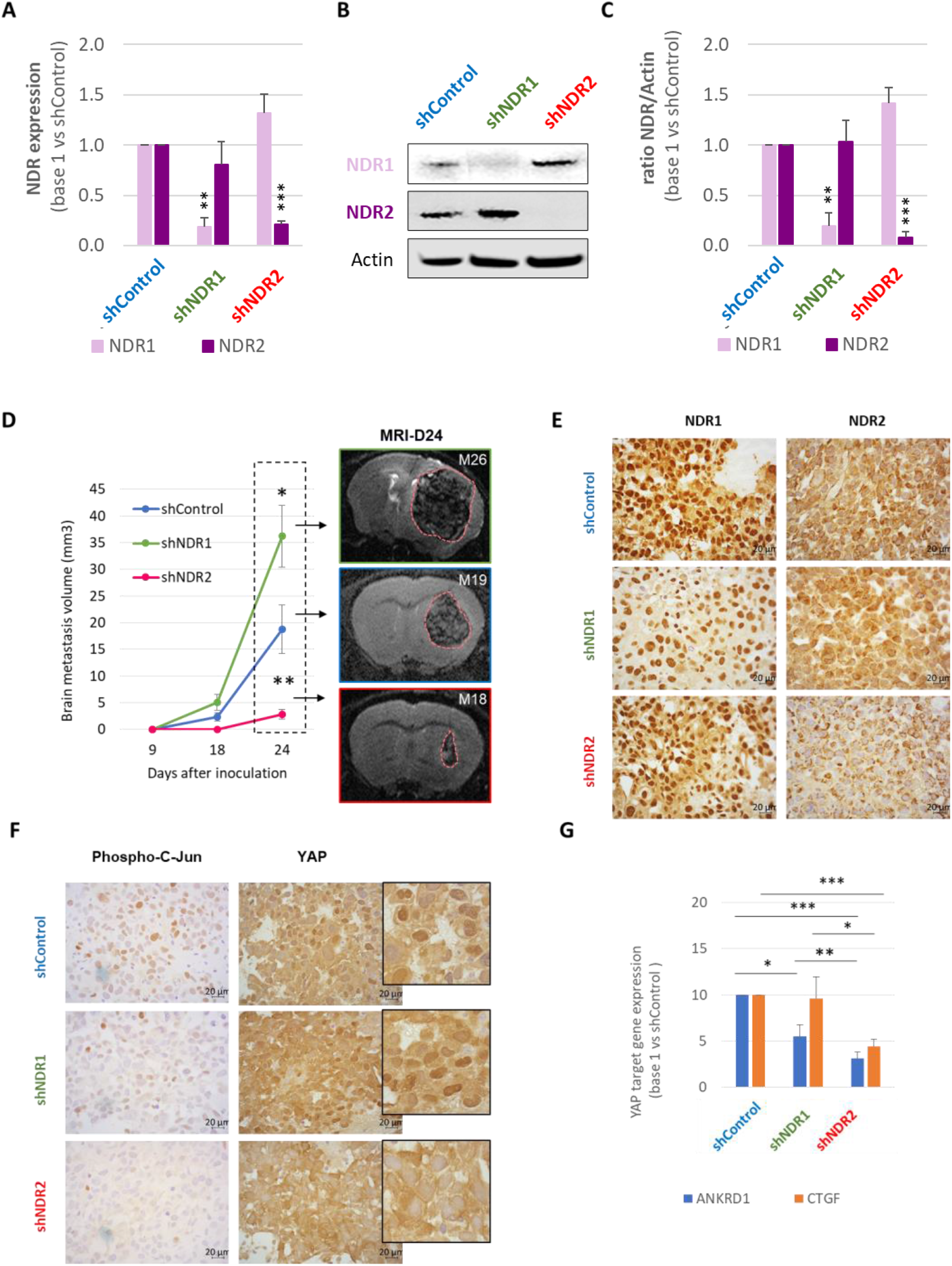
NDR2 silencing reduces development of brain metastases in Swiss Nude. H2030-BrM3 adenocarcinomas of human origin were silenced for NDR1 (shNDR1), NDR2 (shNDR2) or control (shControl) using respective shRNA. **A-C**) Prior *in vivo* experiments silencing was confirmed by RTqPCR (**A**) and western blot (**B, C**) experiments. Mean ± SEM, n=3, **p<0.01 and ***p<0.001 vs respective control. H2030-BrM3 (shNDR1, shNDR2 or shControl) were injected in the left caudate putamen (striatum) in Swiss Nude (N=10 animals per condition). Animals were then followed by anatomical MRI over 24 days ‘period to follow brain metastases development. **D**) Quantitative analyzes of tumor volume at 9, 18 and 24 days after cell inoculation using MRI. Mean ± SEM, n=10 *p<0.05 and **p<0.01 vs Control group. Left panel show a representative T2w-MRI images of the lesions at D24 in the three different experimental groups. **E-G**) Representative immunohistochemically analysis of NDR1 and NDR2 (**E**) or phospho-C-Jun and YAP (**F**) staining in brain metastases from the three experimental groups. **G**) mRNA expression of YAP targets genes ANKRD1 and CTGF in H2030-BrM3 silenced for NDR1 (shNDR1), NDR2 (shNDR2) or control (shControl). Mean ± SEM, n=3 *p<0.05, **p<0.01 and ***p<0.001 vs respective control.

Immunostaining confirmed the lower expression of NDR1 and of NDR2 in BM of respective experimental groups (Figure 6E). Furthermore, we observed a reduction of the nuclear staining of phospho-C-Jun and YAP in BM of shNDR2 group (Figure 6F). In these cells expression of YAP target genes, ANKRD1 and CTGF, were reduced in comparison to the shControl or shNDR1 cell expression (Figure 6G).

### NDR2 is more expressed in metastatic NSCLC than in localized NSCLC while not YAP nor phospho-c-Jun

We assay the tumoural expression of NDR2, YAP and phospho-c-Jun in 25 patients with localized cancer and 20 patients with metastatic cancer (Figure 7A-C). NDR2 is more expressed in tumour of metastatic NSCLC (H-score: 193.2 ± 5.8) than in localized NSCLC (136,4 ± 10,7). There was no difference in the expression of NDR2 between the primary and BM tumours (196.0 ± 1.8) of the same patients with NSCLC.

**Figure 7:**
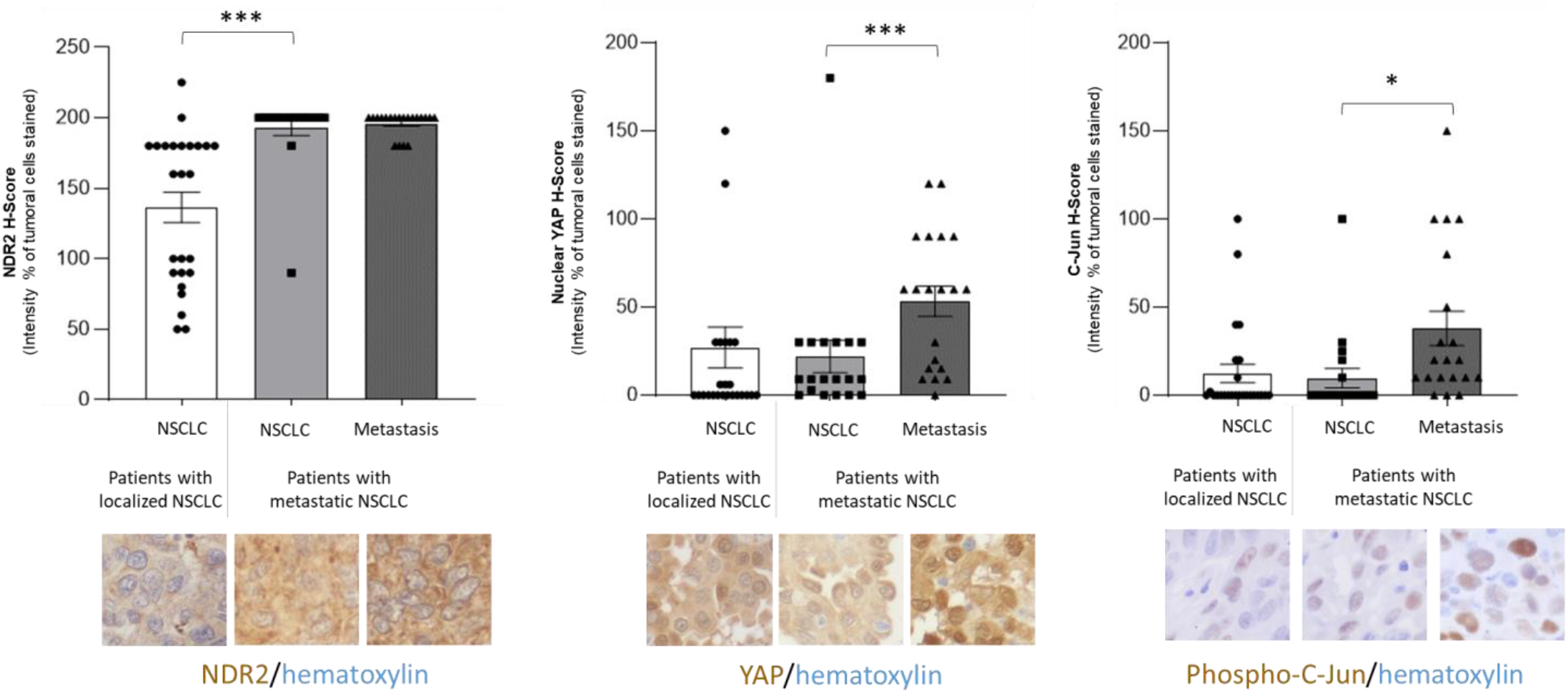
NDR2 expression increases in primitive NSCLC leading to brain metastasis. We selected a retrospective population of 45 patients operated on a non-metastatic NSCLC (n=25) or metastatic NSCLC (n=20) for whom both the primitive tumour and the brain metastasis (BM) were available, at Caen University Hospital After the epitopes retrieval and saturation of endogenous peroxidases, slides with tumour specimen were incubated at room temperature with NDR2 (1:400), YAP (1:400) or phospho-c-jun (1:50) antibody overnight at 4°C. Then, antibody fixation was revealed by the Novolink System (Leica). The staining intensities were evaluated in a blinded manner at 40x magnification and were scored using marker-specific 0–3 scales (0: negative, 1: weak, 2: moderate, and 3: strong). An overall IHC composite score was calculated based on the sum of the staining intensity (0–3) multiplied by the distribution (0%–100%) from all parts of the slide, thereby providing an H-score between 0 and 300. Data are represented as the mean ± SEM. Statistical significance was calculated and p value are indicated by asterisks: *p<0.05.

The nuclear YAP or phospho-c-Jun H-Score are similar between primary tumours of patients with localized NSCLC (YAP: 27.1 ± 11.6, phospho-c-Jun: 12.5 ± 5.3) and those with metastatic NSCLC (YAP: 21.9 ± 9.2, phospho-c-Jun: 9.8 ± 5.4). However, nuclear YAP or phospho-c-Jun are 2 fold higher in BM (YAP: 53.3 ± 8.6, phospho-c-Jun: 38.0 ± 9.7) than in primary tumour of patients with metastatic NSCLC. These results confirm that elevated level of nuclear YAP and phospho-c-Jun are hallmarks of metastatic process acquisition.

## Discussion

Based on the literature, we hypothesized that a hypoxic tumour microenvironment could contribute to the inactivation of the RASSF1A/Hippo pathway during bronchial tumour growth and underlies brain metastases formation. The objectives of this work were therefore to test this hypothesis and to understand by what mechanisms hypoxia acted.

We first confirm that in our hand, human primitive NSCLC as their brain metastases are hypoxic (Ren et al, 2013; Yang et al, 2016; Gao et al, 2017; Corroyer-Dulmont et al., 2021, Zheng et al, 2021; for review: Ancel et al, 2021). We observe that the CAIX H-Score is similar between tumours of patients with localized NSCLC and those with metastatic NSCLC, and comparable between primary tumour and brain metastasis from the same patients (Figure S1). Such result does not mean that the primitive tumours, whether metastatic or not, exhibits similar level of hypoxic, but rather that these tumours are not at the same step of the disease and thus maybe not at hypoxic peak for each.

Then, we assess the ability of human bronchial epithelial cell (HBEC) lines expressing (HBEC-3, BEAS-2B) or not RASSF1A (A549, H1299, H1915, H2030-BRM3) to survive severe hypoxia at 0.2% oxygen (Figures 1, Figure S2, Figure S3). Our results demonstrate that such hypoxia induces apoptosis of HBEC lines but that the latter nevertheless manage to survive this unfavorable environment and even to proliferate, whether or not they express RASSF1A. The fact that HBEC lines used for this work resist such hypoxia is consistent with the work of Polosukhin et al, 2011 having maintained cultures of HBEC in hypoxia (1% oxygen) for up to 28 days in an air-liquid interface.

We then studied the influence of this hypoxia on the functionality of the RASSF1A/Hippo pathway. We discovered that hypoxia caused the inactivation of TAZ in HBEC-3 cells cultured for 48h at 0.2% oxygen but the accumulation of active (dephosphorylated) nuclear YAP in HBEC-3 cells as saw by the transcription of one of its target genes: ANKDR1 (Figure 1). Such results were recovered for the other HBEC lines with a few exceptions for BEAS-2B, H1299 and H1915 cells lines (Figure S4). In the BEAS-2B line, the basal level of YAP expression is high, probably due to their immortalization by SV40, an inhibitor of p53 which leads to the activation of YAP (Mello et al, 2017), and may not increase further. However, the H1299 and H1915 lines also have an inactive p53 mutation but do not exhibit such strong nuclear expression of YAP. Another mechanism that we have not identified must thus account for the absence of variation of YAP in these two lines. The increase in the expression of the target genes of YAP by RT-qPCR shows that this transcriptional co-factor is indeed active in the different lines, even if the level of expression of these different target genes is sometimes low and varies between them. Although the BEAS-2B line shows a strong expression of nuclear YAP, the expression of the mRNAs of the YAP target genes is not higher than in the other cell models studied. However, the western blots confirm that YAP is under its active form since phosphoSer127-YAP (inactive form) is either stabilized or slightly reduced in hypoxia.

That hypoxia act differently on YAP and TAZ was already described in ovarian cancer, in which it is described that in hypoxic condition, the phosphorylation/inactivation of YAP decreases while that of TAZ sharply increases (Yan et al, 2014).

The activation of YAP under severe and prolonged hypoxia is supported by the silencing of its negative regulators: RASSF1A and Hippo kinases but not NDR2 (Figure 1, Figure S5). That hypoxia inhibits Hippo kinases and promotes the nuclear localization of YAP as well as its transcriptional activity had already been reported in breast cancer (Ma et al, 2017), liver (Zhang et al, 2018), the colon (Greenhough et al, 2018), the pancreas (Wei et al. 2017) or the ovary (Yan et al, 2014) but not yet in lung cancer. Since hypoxia could influence the Hippo pathway through epigenetic modifications (Thienpont et al, 2016; Palarkurthy et al, 2009), we determined the methylation status of promoters of genes encoding members of the RASSF1A/Hippo pathway. We show that hypoxia does not induce methylation of the promoters from Hippo kinases or *ANKRD1* genes (ANKRD1 was studied since Jimenez et al, 2017 reported frequent methylation of the promoter of this gene in bronchial tumours and that we observed a very strong transcription of this gene following hypoxia) nor does it demethylate the RASSF1A promoter in A549 cells (Figure S6). The decrease in expression of RASSF1A and of the kinases of the Hippo pathway induced by hypoxia does not therefore imply a modification of the methylation status of the promoters of the genes encoding these proteins, the mechanism of action remains to be determined but could involve ubiquitin-dependent regulations, as described in the breast cancer model in which SIAH2 directs the LATS2 kinase to the proteasome (Ma et al, 2017). Indeed, all the members of the Hippo pathway are subject to regulation by ubiquitinylation (Nguyen and Kugler, 2018). The role of the ubiquitynylation should thus be address in further investigations to understand such downregulation.

We next investigate the consequences of YAP activation by severe and prolonged hypoxia in HBEC cells, since active YAP leads the tumourigenic process in bronchial cells. Indeed, YAP is activated in HBEC cells cultured under hypoxia and is described to transcribe genes involved in TEM and cell movement (Dubois et al, 2016). We evaluated the effect of hypoxia on the cytoarchitecture of HBEC-3 cells, their TEM and their 2D motility (Figure 3). The immunostaining of the actin and tubulin filaments that we carried out first of all confirmed that the inhibition of the expression of RASSF1A alters the morphology of the cells which become either very large or very stretched in normoxia (Dubois et al, 2016). Here, we report in an original way that the alteration in cell morphology is enhanced when cells are grown under severe hypoxia. Again, that hypoxia alters the morphology of bronchial cells is in agreement with the work of Polosukhin et al, 2011 who reports that hypoxia affects the differentiation of HBEC *in vitro*: these HBEC cells cultured at an air-liquid interface do not more succeed in forming cilia at their apex, and adopt a mucoid phenotype. This change in morphology is in agreement with **i)** the TEM that we report in parallel in these cells, **ii)** the fact that they undo their cell junctions (adherent and communicating) and **iii)** the fact that they strongly express fascine, molecule known for its involvement in the formation of filopodia (fine cytoplasmic extensions) but also in the extensibility of cells (Tanaka et al, 2019) as well as cell migration (Liang et al, 2016). This morphological/phenotypic change explains why human bronchial epithelial cells adopt an individual type of migration when they are cultured in hypoxia and when they are brought to fill a mechanical wound made on their cell layer while in normoxia, their migration is collective. The hypoxia-induced amoeboid migration has only been described once to date and to our knowledge in a head and neck cancer model and is linked to HIF-1A (Lehmann et al, 2017). We also observe that the HBEC grown in hypoxia do not efficiently repair the wound: HBEC move faster in hypoxia than in normoxia, but do not only migrate toward the other bank, particularly RASSF1A-depleted HBEC. This disorganized migration could be explained by the fact HBEC-3 cells grown in hypoxia and depleted for RASSF1A that strongly express the fascine which, as mentioned above, controls cell movement and elasticity. An increase in fascin has already been described in many cancers, in particular in NSCLC, where it predicts a poorer prognosis (Zhang et al, 2018) because it promotes cell migration and invasion of NSCLC (Zhao et al, 2018). An increase in fascin has also already been described when cells are in hypoxia (Zhao et al, 2014). However, fascin is not a suitable therapeutic target since we observed that its inhibition by siRNA caused major cytonuclear abnormalities in HBEC (data not shown).

We know that YAP is responsible for the increased collective migration rate induced by the loss of RASSF1A expression in HBEC cultured in normoxia (Dubois et al, 2016). We show here that, the gain in migration velocity induced by the RASSF1A depletion independent of YAP but could be dependent of HIF-1A which is stabilized by the loss of RASSF1A and/or YAP expression when HBEC are grown in hypoxia. This result is unexpected since, it was shown that RASSF1A stabilized HIF-1A in NSCLC cells (Dabral et al, 2019) or that YAP stabilized HIF-1A (Zhang et al, 2018). It is therefore probable that the mechanisms allowing the stabilization of HIF-1A in the absence of RASSF1A or YAP are different, and could for example involve the transcription factor ETS-1 (v-ets erythroblastosis virus E26 oncogene homolog 1), which governs the gene expression of HIF-1A (Salnikow et al, 2008) and is itself activated by JNK signaling (Zhang et al, 2013), which is repressed by RASSF1A (Whang et al, 2005). It should also be noted that the experiments that we carried out in hypoxia were done in percentage of O2 and for a duration different from the work of Dabral, which can also account for the differences observed between our respective studies and suggest that the mechanisms are specific to a level of hypoxia (moderate, severe, chronic, etc.).

As previously mentioned, the YAP activity in hypoxia coincides with the loss of expression of the Hippo pathway kinases in hypoxia in all lines except the NDR2 kinase, which appears to be stabilized in hypoxia. This stabilization of NDR2 in hypoxia is therefore consistent with the results obtained previously showing that hypoxia promotes the nuclear localization of YAP in certain lines of NSCLC and the expression of the target genes of YAP. Indeed, this kinase can, through the GTPase RhoB pathway induce the nuclear translocation of YAP (Keller et al, 2019). It would then be interesting to study the Rho protein pathway within the different lines in order to better understand by which mechanisms the NDR2 kinase promotes the nuclear localization of YAP and therefore its activation in our different NSCLC lines. Hypoxia has also been shown to induce cell migration of cancer cells via the RhoA pathway (Leong & Chambers, 2014).

The role of the NDR2 kinase and the relationship with hypoxia having been investigated *in vitr*o on cells derived from NSCLC, we subsequently studied the expression of this kinase and of YAP on tumour samples from affected patients of CBNPC and followed at the CHU of Caen (Figure 7). On these samples, we performed immunohistochemistry in order to quantify the expression of NDR2, YAP and CAIX as a marker of hypoxia (Ambrosio et al, 2016). We show that NDR2 kinase expression is greater in tumours from patients with metastatic NSCLC suggesting a link between the metastatic process and NDR2 expression. In addition, the YAP cofactor is more often expressed in primary tumours of patients with metastatic NSCLC than in patients with non-metastatic NSCLC. These observations highlight its potential role in the development of metastases, by inducing the expression of pro-metastatic target genes as has already been shown in breast cancer and melanoma cells (Lamar et al, 2019). The results also show that the expression of YAP at the nuclear level is greater in the metastasis compared to that found in the primary tumour, which again suggests that the development of these metastases takes place in a YAP-dependent manner although this remains to be proven on an in-vivo study model by carrying out an extinction of the expression of YAP or else by preventing its nuclear translocation.

Regarding the expression of YAP, the results did not reveal a significant difference in expression between the metastatic or non-metastatic NSCLC. Such comparison was not previously done by others to our knowledge but does not call into question the role of YAP in the formation of carcinoma metastases, in particular from lung, since independently of the quantity of YAP, the important thing is its activation.

We finally used a model of lung cancer derived BM in mice to test the NDR2 impact on formation of such xenograft and growth (Figure 6, Figure S7). This model shows that silencing NDR2 kinase (but not NDR1) reduces the number of metastases and the overall volume and rate of lesion progression. Collectively, these results are therefore in favor of an effect of metastatic promotion of NDR2 and consistent with the role of the kinase NDR2 involved in the control of cell movement via the regulation of YAP demonstrating a pro-metastatic effect of the latter (Dubois et al, 2016).

## Conclusion

In conclusion, our results demonstrate that hypoxia is an aggravating factor in bronchial carcinogenesis by influencing the RASSF1A/Hippo pathway in bronchial cell lines. In fact, hypoxia plays a role in cell viability but also in morphological changes in cells with a more invasive and amoeboid phenotype. This hypoxia affects cell migration, the cells opt for individual rather than collective migration, an effect all the more marked when RASSF1A is inhibited. These changes appear to be related to the fact that hypoxia induces inappropriate activation of YAP, probably by modulating regulators and kinases of the Hippo pathway and not by epigenetic modifications.

These new data improve our understanding of the relationship between the tumour microenvironment, the Hippo signaling pathway and the adaptation of bronchial tumour cells. Our results shed light on HIF1 as a potential therapeutic target in patients with NSCLC with inactivation of the RASSF1A gene, but also YAP. Thus, the pharmacological targeting of these new targets could be effective in preventing the spread of cancer and in improving the vital prognosis of patients with NSCLC. Our results also indicated that NDR2 kinase is over-active in NSCLC in part by hypoxia and supports BM formation. NDR2 expression is thus a useful biomarker to predict the metastases risk in patients with NSCLC, easily measurable routinely by immunohistochemistry on tumour specimens

**Table 1.** NDR2 silencing decreases formation and growth rate of BM from HBEC.

|  | H2030-BrM3 |  |  | A549 |  |
| --- | --- | --- | --- | --- | --- |
|  | shControl | shNDR1 | shNDR2 | shControl | shNDR2 |
| Animals with BM<br>n (%) | 7/10<br>(70%) | 7/10<br>(70%) | 6/10<br>(60%) | 8/10<br>(80%) | 5/9<br>(55.5%) |
| BM location |  |  |  |  |  |
| *Striatum* | 7/7 (100%) | 7/7 (100%) | 6/7 (85.7%) | 8/8 (100%) | 0 |
| *Cortex* | 0 | 2/7 (28.6%) | 0 | 1/8 (12.5%) | 5/5 (100%) |
| Start growing<br>(day after inoculation) | 19.71±1.11 | 18.00±0.00 | 24.00±0.00* | 33.0±6.3 | 40.6±5.5* |
| Growth rate<br>(mm<sup>3</sup>/day) | 18.73±4.53 | 36.22±5.81* | 2.84±0.90** , ### | 7.27±1.57 | 0.33±0.13** |

t-test : \* vs shControl ; # vs shNDR1

## Additional Information

### Ethics approval and consent to participate

The study was conducted according to the guidelines of the Declaration of Helsinki, and approved by the Institutional Ethics Committee of the Caen University Hospital center (protocol code DC-2008-588, date of approval: 12th April 2010) authorizing the collection, conservation and preparation activities for scientific purposes included in this collection of human biological samples at the Caen University Hospital center.”. Informed consent was obtained from all subjects involved in the study.

### Availability of data and material

All data are stored at the GIP Cyceron and Université de Caen Normandie (Caen, France), and can be made available upon request.

### Authors’ contributions

`o` Conception and design: GL, JL, EB.
`o` Development of methodology: JL, TB, CB, MD, FD, DLF, JT, ST, JT, MB, SV, GL.
`o` Acquisition of data (provided animals, acquired and managed patients, provided facilities, etc.): JL, TB, CB, MD, FD, DLF, JT, ST, JT, MB, SV, EB, GL.
`o` Analysis and interpretation of data (e.g., statistical analysis, biostatistics,computational analysis): JL, TB, CB, MD, FD, DLF, JT, ST, SV, EB, GL.
`o` Writing, review, and/or revision of the manuscript: JL, TB, CB, MD, FD, DLF, JT, ST, JT, MB, SV, EB, GL.
`o` Administrative, technical, or material support (i.e., reporting or organizing data, constructing databases): JL, TB, CB, MD, FD, DLF, JT, ST, SV, GL.
`o` Study supervision: GL, EB.

## Acknowledgments

Authors thanks the VIRTUAL’HIS platform. We also thank Dr. Joan Massagué (MSKCC, USA) for providing the H2030-Br3M cell line.

